# KinaFrag explores the kinase-ligand fragment interaction space for selective kinase inhibitor discovery

**DOI:** 10.1101/2021.03.28.436965

**Authors:** Zhi-Zheng Wang, Xing-Xing Shi, Fan Wang, Ge-Fei Hao, Guang-Fu Yang

## Abstract

Protein kinases play a crucial role in many cellular signaling processes, making them one of the most important families of drug targets. But selectivity put a barrier at the design of kinase inhibitors. Fragment-based drug design strategies have been successfully applied to develop novel selective kinase inhibitors. However, the complicate kinase-fragment interaction and fragment-to-lead process pose challenges to fragment-based kinase discovery. Here, we developed a web source KinaFrag to investigate kinase-fragment interaction space and perform fragment-to-lead optimization. KinaFrag contained 31464 fragments from reported kinase inhibitors, which involved 3244 crystal fragment structures and 7783 crystal kinase-fragment complexes. These crystal fragments were classified by their binding cleft and subpockets, and their 3D structure and interactions were displayed in KinaFrag. In addition, the structural information, physicochemical information, similarity information, and substructure relationship information were contained in KinaFrag. Moreover, a computational fragment growing strategy obviously developed by our group was implemented in the KinaFrag. We test this fragment growing strategy using our fragment libraries, and obtained a lead compound of c-Met with ~1000-fold *in vitro* activity improvement compared with the hit compound. We hope KinaFrag could become a powerful tool for the fragment-based kinase inhibitor design. KinaFrag is freely available at http://chemyang.ccnu.edu.cn/ccb/database/KinaFrag/.

**Biographical note:** Zhi-Zheng Wang is a PhD student at College of Chemistry, Central China Normal University (CCNU), and the direction of his thesis is computational molecular simulation.

Xing-Xing Shi is PhD student at College of Chemistry, of CCNU, and the direction of her thesis is computational molecular simulation.

Fan Wang is a lecturer at College of Chemistry of CCNU. He received the PhD degree in Computational Chemistry from University of Amiens, France.

Ge-Fei Hao is Professor in Bioinformatics in College of Chemistry of CCNU. He received his PhD in Pesticide Science from CCNU.

Guang-Fu Yang is Professor in Chemical Biology. He is the group leader and has been the Dean at College of Chemistry of CCNU. He received the PhD degree in Pesticide Science from Nankai University, Tianjin, China.

## Introduction

Protein kinases (PKs) play vital roles in the signal transduction pathways, therefore have been regarded as important drug targets for multiple diseases in the past two decades.^1,2^ A series of efforts have been paid to the discovery of small-molecule protein kinase inhibitors (PKIs), which have achieved impressive success in the therapy of several serious diseases, including leukemia, cancers and infectious diseases. Up to now, 62 small-molecule PKI drugs are approved into market by the US Food and Drug Administration (FDA) according to the Blue Ridge Institute for Medical Research in Horse Shoe (http://www.brimr.org/PKI/PKIs.htm) and PKIDB. Moreover, the number of approved and clinical PKIs are increasing over the years. However, most of the reported PKIs bind into the ATP active site of kinases. The kinase selectivity has emerged as a significant problem for the development of PKIs.^3^ Hence, the design of highly selective PKIs is still a major challenge and a clinical necessity.

Fragment-based drug design (FBDD) has become a widely used drug discovery strategy to decease the time cost.^4^ Fragments usually refers to the molecules with low molecular weight (MW) and complexity, FBDD is ability to explore bigger chemical spaces. In addition, FBDD can achieve higher hit rates in screening. It is easy to design compounds with favorable physicochemical properties, high ligand efficiency (LE), and chemical novelty using FBDD. However, the small in size of fragments also has some disadvantages: (1) The generation of fragment database for wider chemical space. (2) The detailed interaction between target and fragment is hard to be obtained because of the weak binding affinity. (3) the lack of fragment binding mode poses a challenge to the fragment evolution.^5,6^ But according to a recent study, fragments usually keep in a conserved conformation during the optimization. Consequently, the understanding of known protein-fragment interactions is important for the whole FBDD process.

Protein kinases are regarded as one of the most commonly applied targets for FBDD.^7^ There are a series of PKIs discovered through FBDD. The ligand binding hot spots, hinge region residues, could be used as primary fragment hit binding site, which is more likely to achieve high activity in fragment screening. Meanwhile, the abundant reported PKIs and kinase-fragment interaction information can be utilized as guidance for fragment optimization. During the fragment to lead process, FBDD could meet the requirement of selective PKIs by maximizing the interactions with kinase subpockets. These advantages lead to an efficient discovery of selective PKIs via FBDD. However, how to realize the kinase-fragment interactions and used proper fragments to design highly selective PKIs are still big challenges.

In order to facilitate the fragment-based selective kinase inhibitor discovery, we construct a comprehensive and free available web source KinaFrag (http://chemyang.ccnu.edu.cn/ccb/database/KinaFrag/. KinaFrag could investigate the reported kinase bioactive chemical structures and explore the kinase-fragment interaction space. In addition, a fragment growing strategy was implemented for the exploration of unknown kinase-fragment interaction space. A c-Met lead compound was discovered with a more than 1000-fold activity improvement using KinaFrag. We hope KinaFrag could be a powerful tool for the fragment-based kinase inhibitor discovery, structure-based drug discovery and prediction model generation (**Figure 1**).

**Figure 1.**
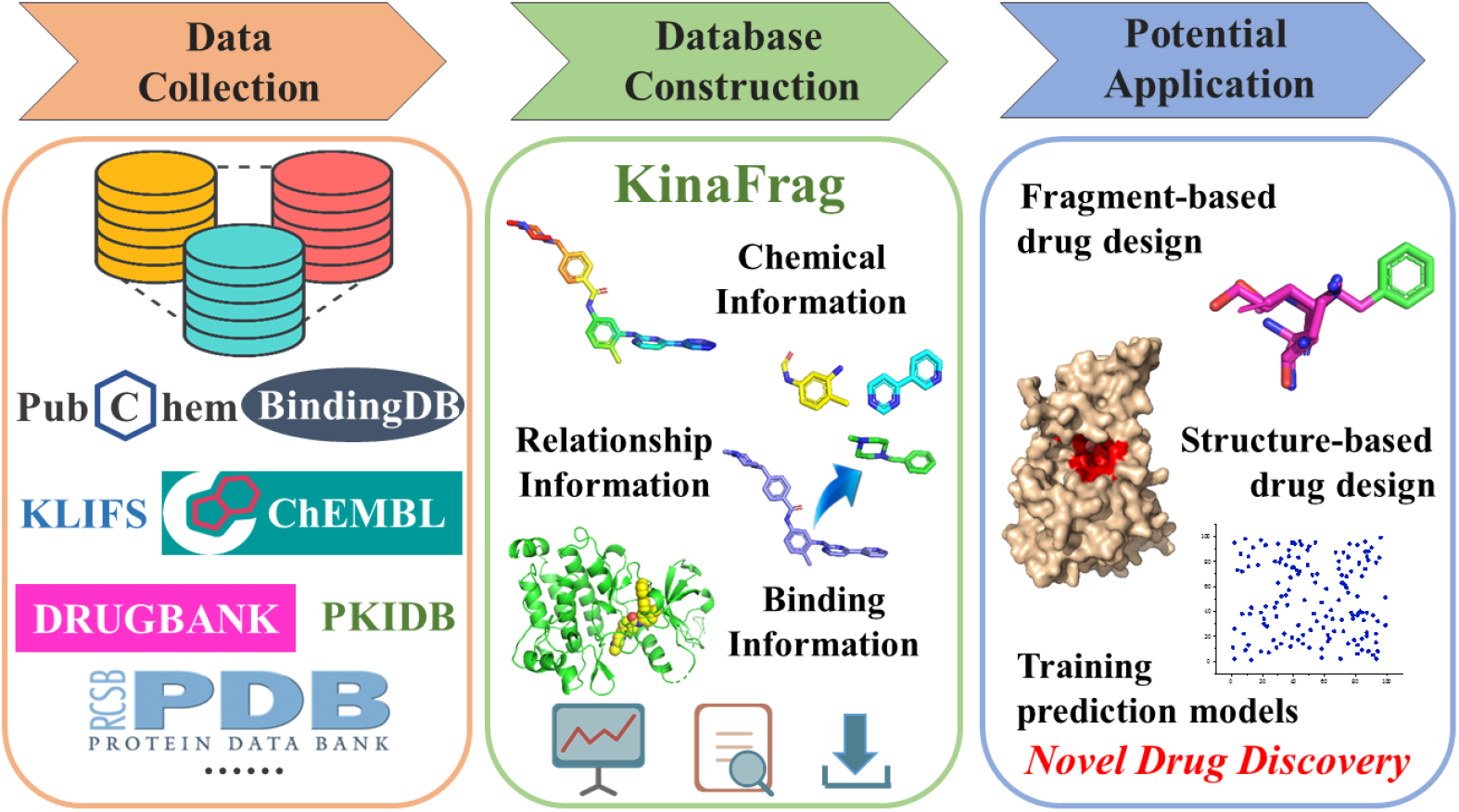
The construction of KinaFrag.

## Materials and methods

### Data collection

The kinase inhibitors used for the fragment generation in KinaFrag was collected from scientific literatures and several commonly used public databases. The kinase list was extract from Kinase.com (http://www.kinase.com/web/current/), which contained 518 human kinases. All the crystal structures of kinase-inhibitor complexes were retrieved from KLIFS database (https://klifs.net/).{Kooistra, 2016 #8} Meanwhile, more than 100,000 reported kinase inhibitors were collected from more than 5000 literatures and some common databases, e. g. PubChem,^8^ BindingDB,^9^ DRUGBANK,^10^ ChEMBL.^11^ To achieve better bioactive chemical space, these compounds were filter by bioactivity lower than 1 μM (IC_50_, Ki, EC_50_, or KD_50_). Although from authoritative web source, there were still some mistakes with obtained compounds, therefore these compounds would be excluded from our further study. For the remained kinase inhibitors, the replicate compounds were combined. The substrates and substrate derivative ligands (such as ATP, ADP) and covalent inhibitors would be discarded.

### Fragmentation and classification

The active protein kinase domain exhibits similar sequence and structural futures. It can be divided into front cleft (usually refers to the ATP binding site), gate area, and back cleft.^6^ These binding pockets are mainly featured by three key important structures of the kinase: aspartate-phenylalanine-glycine (DFG) motif, αC-helix, and gatekeeper (GK) residue adjacent to the hinge region. Most of current PKIs bind in the adenine pocket (AP), which is located in the front cleft, and form hydrogen bonds with hinge region. Meanwhile, there are three other subpockets in the front cleft: the solvent exposed loop (linker subpocket), front pocket I (FP-I) and front pocket II (FP-II). In the gate area and back cleft, multiple subpockets exist (BP-I, BP-II, BP-III, BP-IV, BP-V), and the subpocket size depending on the conformation of DFG motif and αC-helix.^5^ On the basis of these clefts and subpockets, PKIs could be classified in different types.

The generation of fragments is important to the build of fragment database. In KinaFrag, the kinase inhibitor fragmentation steps were similar to our previous study. All the inhibitors were first cut into fragments according to the rotatable bonds using DAIM software. For the fragments generated from crystal structure, they were classified by the binding cleft and subpockets according to KLIFS, and all generated fragments would be remained. For the fragments obtained from non-crystal PKIs, they were filter by the fragment-like “rule of three” as: MW < 300, number of hydrogen bond acceptor ⩽ 3; and cLogP ⩽ 3 (the logP of a compound is a measure of the compounds’ hydrophilicity, where P is the partition coefficient between n-octanol and water log[Coctanol/Cwater]). Only fragments with proper physicochemical properties are retained. In addition, the duplicate fragments were dropped. During the fragmentation, the linking point in the generated fragments were remained.

### Data reorganization

After the fragment generation, all the fragments used for our database would be reorganized. The structural information of fragments was obtained with Openbabel^12^ and RDKit, including formula, canonical smiles, InChI, InChI Key and MACCS Fingerprint, which would be convenient for the structure search. Meanwhile, a series of important physicochemical properties besides “rule of three” were calculated with RDKit and ALOGPS to explore the chemical space of kinase binding fragments, such as number of heavy atoms, clogS and number of rings. The substructure relationship analysis between inhibitors and fragments were recorded to explore the potent fragment binding target. The similarity between each fragment was also calculated as the fragment replacement was often used in FBDD. The target information of fragments was collected since the fragment binding exhibit an obvious conservation which might be helpful to identify fragments with similar binding position. Moreover, for the fragments from crystal structures, their binding modes with kinases were systematically analyzed. Their binding affinity towards binding kinase were calculated with Molecular Mechanics Poisson Boltzmann Surface Area (MM/PBSA) method, and the ligand efficiency (LE) value was also computed. LE value was the measurement of ligand optimization potential, and could be defined as the ratio of Gibbs free energy (ΔG) to the number of non-hydrogen atoms of the fragment. The interactions between kinase and fragments were described using PLIP^13^ (https://projects.biotec.tu-dresden.de/plip-web/plip/index) and LigPlot,^14^ where hydrophobic interaction, hydrogen bond, π-π stacking, halogen bond, salt bridge and metal complexation could be analyzed.

### Fragment-growing workflow

The optimization from fragment hit to lead compounds was a significant challenge for fragment-based drug discovery. In our previous study, we developed a fragment growing protocol named pharmacophore-linked fragment virtual screening (PFVS) for the discovery of *bc1* inhibitors.^15^ And we developed the world first fragment-based drug discovery web server Auto Computational Fragment *in silico* Screening (ACFIS).^16^ In KinaFrag, the fragment-growing strategy in ACFIS was implemented for fragment-based kinase inhibitor design. Users are required to provide a kinase-fragment complex first, and this fragment would be regard as starting structure for fragment growing. If the complex was come from KinaFrag, the linking point was determined, and if the complex was upload by users, the linking point need to be specified. Then at least one database of subpocket binding fragments could be selected for compounds generation. The number of output compounds, job name, e-mail address and password were optional for the task and viewing result. Once the parameters were set, fragments in database were linking with starting structure. Amber16 was employed to perform an energy minimization and a short-time molecular dynamics simulation, while general amber force field (gaff) was used to generate topology and parameter files for compounds. The binding free energy between generated compounds and kinase would be calculated with MM/PBSA method. The generated compounds, binding free energy, physicochemical properties and their binding modes would be exhibited in the web page.

### Free energy calculation

The binding free energy (Δ*G_binding_*) was calculated by the combination of MM/PBSA method for the enthalpy and an empirical method for the entropy as following formula:^17^

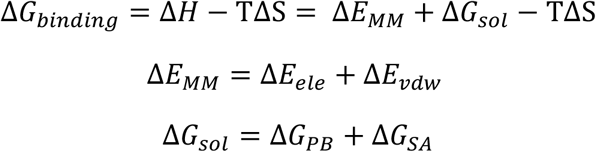

In which Δ*G_binding_* was the binding free energy of ligand, Δ*H* and –TΔS were the enthalpy and the entropy, respectively. The changes in the gas phase molecular mechanics (MM) energy Δ*E_MM_* could be calculated though electrostatic energies Δ*E_ele_*, and the van der Waals energies Δ*E_vdw_*. While solvation free energy Δ*G_sol_* is the sum of the electrostatic solvation energy Δ*G_PB_* (polar contribution) and the nonpolar contribution Δ*G_SA_* between the solute and the continuum solvent. The polar contribution is calculated using the PB model, while the nonpolar energy is usually estimated using the solvent-accessible surface area (SASA).

### Web site configuration

The web front end of KinaFrag was constructed by using PHP (version 5.3), which was run by Apache HTTP Server (version 2.2.21) on the Linux computer cluster. To be specific, the information and results of each task are stored in a MySQL database (version 14.12). Visualization of molecular structure was provided by JSmol (http://www.jmol.org/) apple. IE10 (or later version), Firefox and Google Chrome are recommended browsers for KinaFrag web with the best view resolution of 1024 × 900. The fragment-growing workflow were developed by using Python2.7, which were manipulated to run by Torque-6.0.1 and Maui-3.3.1.

## Results and discussion

### Data statistics in KinaFrag

After data preparation and reorganization, KinaFrag were built freely for all users (**Figure 2A**). In statistics, KinaFrag contains 31464 kinase ligand fragments, which were generated from 3166 crystal kinase inhibitors and 80039 non-crystal structures. Among the 31464 fragments, 3244 unique fragments were come from crystal structures, which involved 4218 PDB structures. Finally, 7783 kinase-fragment complexes were obtained. For the specified binding site, front cleft took up most of the binding fragments. There were 1804 kinase-fragment complexes of Linker subpocket (**Figure 2B**), and 799 fragments were unique (**Table 1**). At the AP subpocket, 4003 kinase-fragment complexes were obtained, while consist of 1290 non-duplicated fragments. There were 996 kinase-fragment complexes of FP pocket (FP-I and FP-II), and 409 fragments were exclusive. These results coincided with the current situation that front cleft binding inhibitors occupied around 95% of all PKIs. Besides front cleft, 589 kinase-fragment complexes of gate area (BP-I-A sand BP-I-B) were identified which contained 372 unique fragments. The gate cleft was a linker between front and back cleft, and shared a conserved sequence, which lead to a limited number of binding fragments. There were 391 kinase-fragment complexes of back cleft (BP-II, BP-III, BP-IV, BP-V), and 297 fragments were unique. The low number of gate area and back cleft bound fragments indicated that inhibitors could target these regions might need more attention.

**Figure 2.**
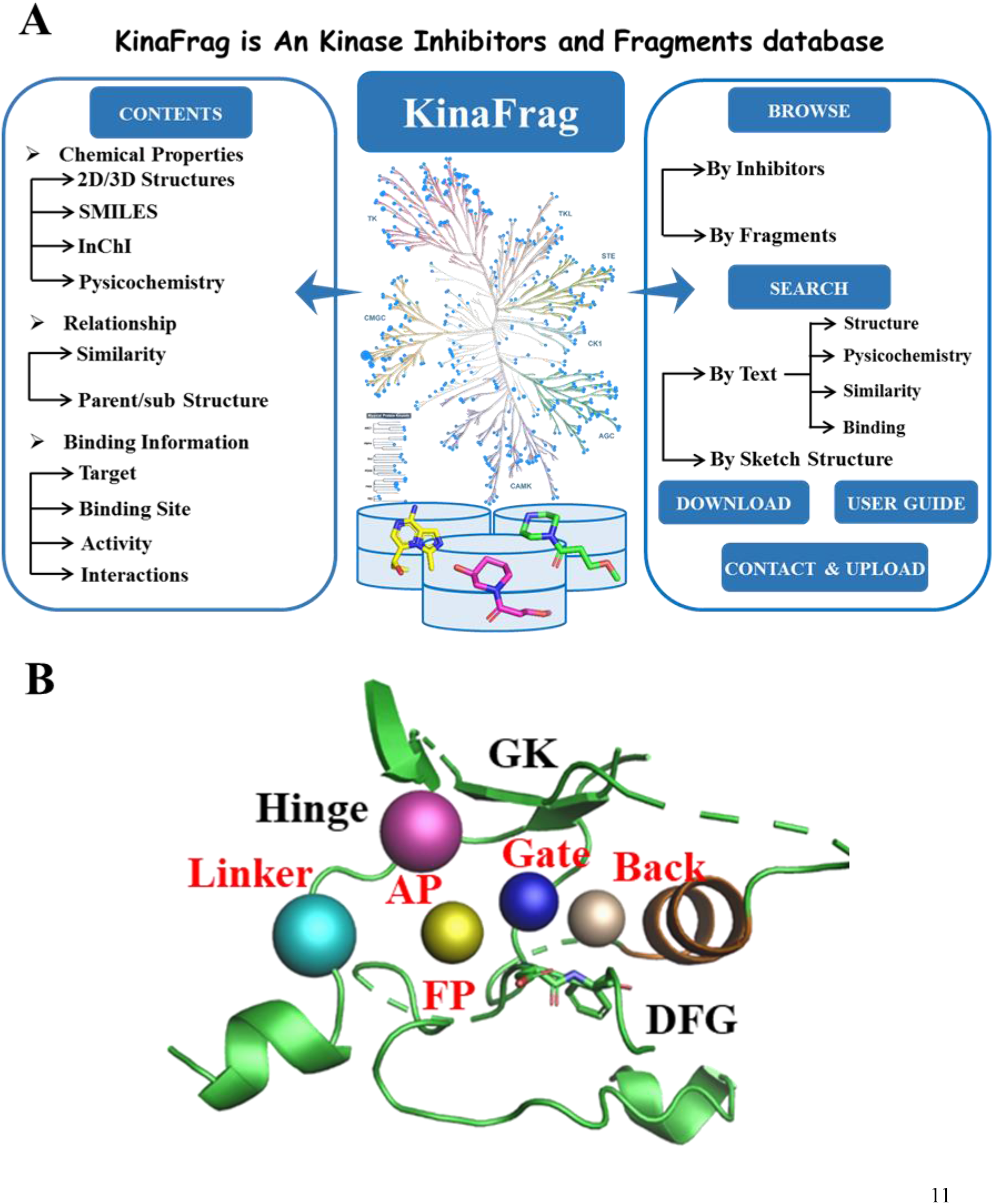
The content of KinaFrag. (A) The chemical property, relationship and binding information of kinase binding fragments were included in KinaFrag. (B) The binding pocket divided in KinaFrag.

**Table 1.**
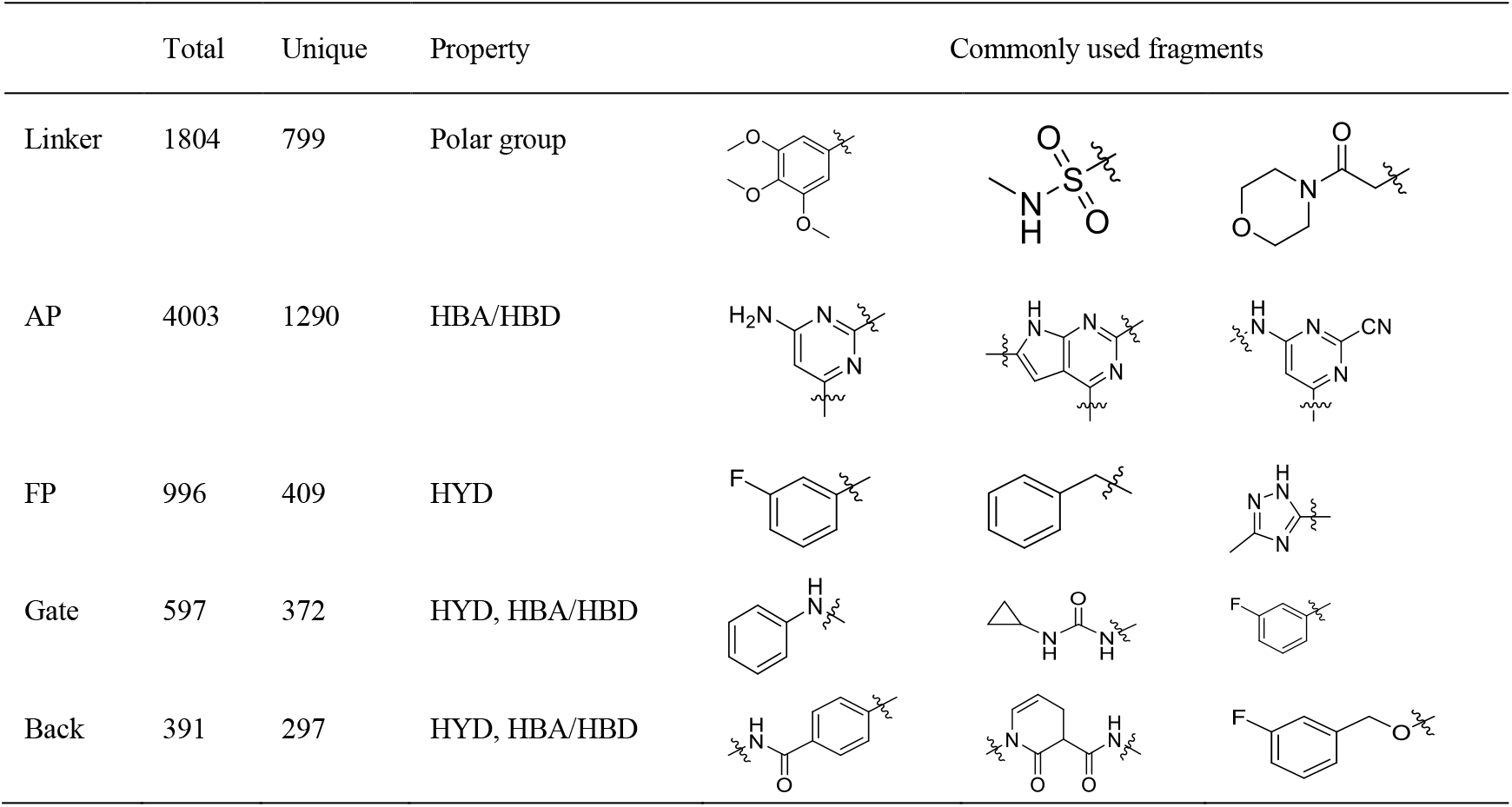
Properties of binding fragments of KinaFrag.

|  | Total | Unique | Property | Commonly used fragments |
| --- | --- | --- | --- | --- |
| Linker | 1804 | 799 | Polar group |  |
| AP | 4003 | 1290 | HBA/HBD |  |
| FP | 996 | 409 | HYD |  |
| Gate | 597 | 372 | HYD, HBA/HBD |  |
| Back | 391 | 297 | HYD, HBA/HBD |  |

We summarized the size and properties of subpocket bound fragments (**Table 1**). The Linker bound fragments usually exposed and interact with solvent, and are usually polar groups. For fragments located in the AP subpocket, they always form hydrogen bounds with the hinge region residues. Therefore, they contained hydrogen bond acceptor (HBA) and/or hydrogen bond donor (HBD). The FP is a hydrophobic area, and fragment bound there are often hydrophobic groups (HYD). The gate area is another hydrophobic region, but some PKIs form hydrogen bonds with DFG motif. And fragments often contain amide groups. The back cleft is a hydrophobic cavity, but some region is exposed in the solvent. Therefore, fragments bound there exhibited multiple types. During the fragment evolution process, it is important to select suitable subpocket binding fragment to obtain good activity and promising selectivity.

### Fragment information in KinaFrag

The browsing tools summarize the data in KinaFrag through two categories: ‘Browse Inhibitors’ and ‘Browse Fragments’. In the ‘Fragments’ page, all the fragments generated by KinaFrag were listed with their brief introduction, such as formula, structure, canonical SMILES, MW, binding clefts and binding subpockets. The location of kinase clefts and subpockets were put in the top of page. Users could sort all fragments according to their demand. If users were interested with one fragment, they could click the ID to look through the detailed information, which contained six parts (**Figure 3**).

**Figure 3.**
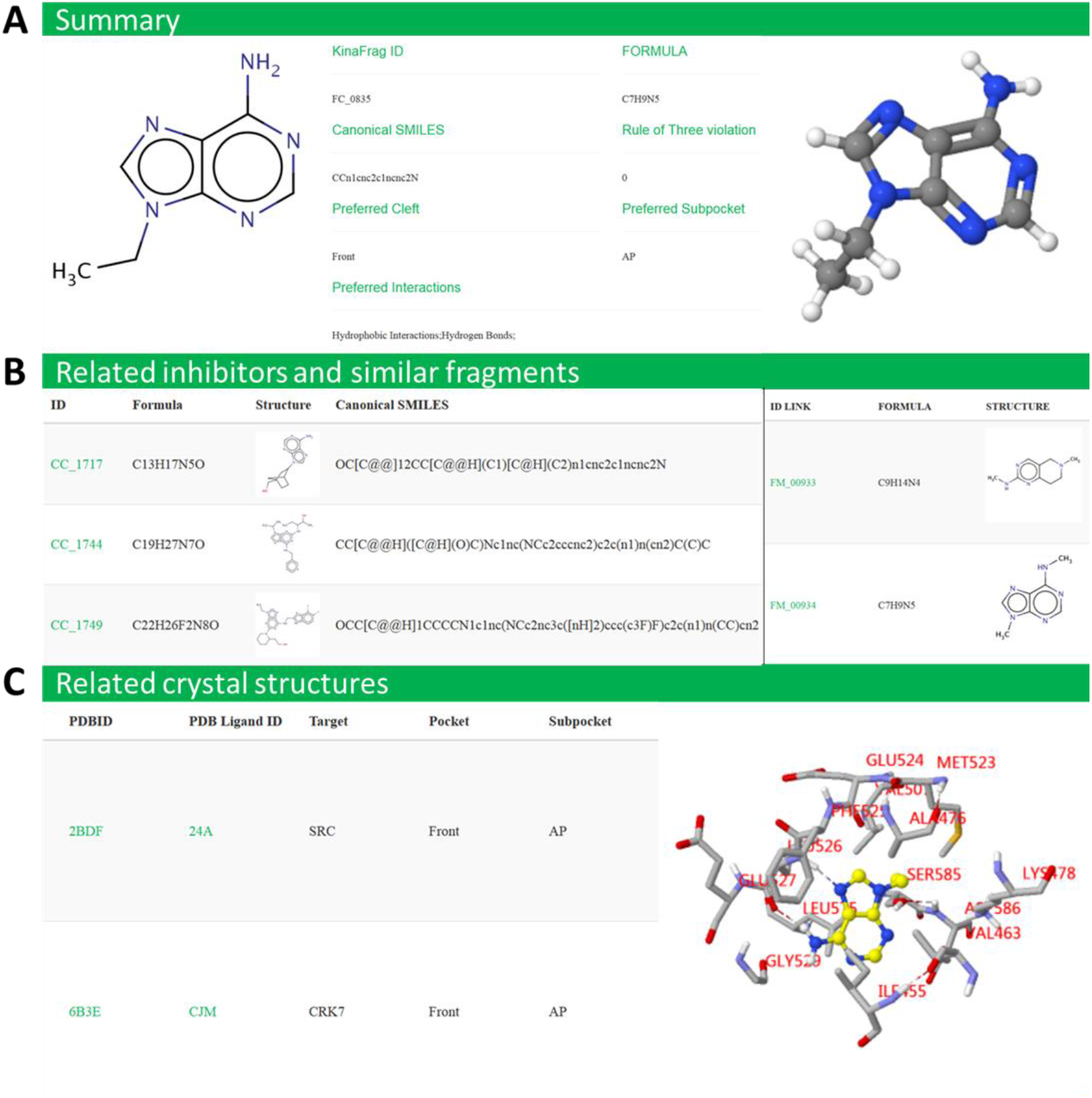
The fragment information in KinaFrag. (A) For a fragment, its structural information, physicochemical information, and binding information were given in summary page. For example, fragment numbered FC_0815 was a AP subpocket binding fragment and usually form hydrogen bond with kinase. (B) The information of related inhibitors that fragment come from, and the similar fragments. (C) The information of fragment binding kinases. Their detailed interaction were showed by JSmol.

In Summary page, the most important properties were introduced, including formula, canonical SMILES, rule of three violation, preferred Cleft, preferred Subpocket and preferred interactions in their original structures. For the Structure page, formula, canonical SMILES, InChI, InChI Key, MACCS fingerprint and 3D conformation were provided. Nine important physicochemical were exhibited in Physicochemical page, including MW, clogP, clogS, number of heavy atoms, number of HBond acceptor, number of HBond donor, total polar surface area, number of rings and number of rotatable bonds, which were important to describe the complexity and hydrophobicity of fragments. In the Drug page, all the parent inhibitors were shown to make users further understand the usage of fragments. For the Similar fragment page, fragments with similarity > 0.5 were given for fragment optimization. As mentioned above, the fragment binding information was the most significant for the FBDD, therefore in Map2PDB page, all the crystal structure contained viewed fragment were exhibited along with their target name, information in RCSB PDB database, and binding location. The detailed interaction information was provided in an individual page. The detailed interaction information contained more specified data, such as the information of target kinase, conformation stage (DFG and αC), binding free energy, 2D and 3D kinase-fragment interaction, and interaction details (residue number and interaction type). As for the collected kinase inhibitors, they shared the similar exhibited way with fragments. All the generated fragments and their original inhibitor could be download freely.

### Query fragments in KinaFrag

In order to facilitate the retrieval of the data in KinaFrag, we provide the searching and browsing tools. As to the searching tools, KinaFrag can be queried with text-based and structure-based search. Text-based search serves as a simple way to search whole database by entering a single term. In KinaFrag, multiple keywords could be used for fragment search, such as SMILES, InChI, binding cleft, binding subpockets and interaction type. After search demand was imported, all the compounds that meet the requirements would be displayed. For structure-based search, users can sketch a molecule within the JSME editor. When the imported structure was submitted, the queried fragments and similar would be exhibited and sorted by similarity. And the brief description, such as formula, structure, canonical SMILES, MW, binding clefts and binding subpockets were also displayed. For the search result, all the text-format information like structural information, target information and binding information as well as 3D structures in MOL2/PDB format could be download.

**Figure 4.**
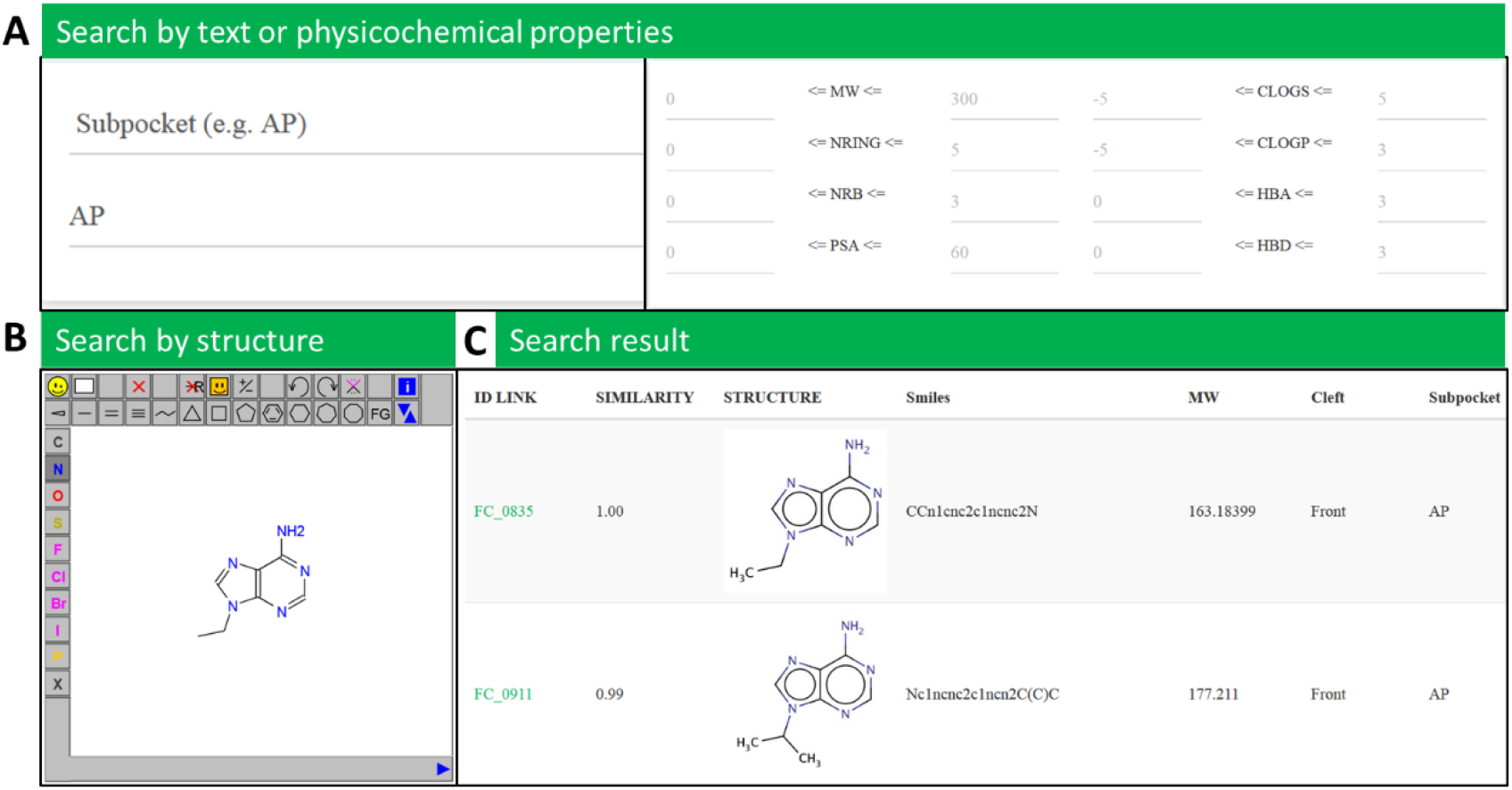
The search function in KinaFrag. (A) Fragments in KinaFrag could be searched by text, including formula, Smiles, InChI, InChIkey, binding cleft, subpockets and interaction type with target, or by physicochemical properties. (B) Users could search queried fragment with chemical structure. (C) The result fragments were list after the search, and the fragment was sorted by the similarity with queried fragment.

### Case study: design potent c-Met inhibitors with KinaFrag

The receptor tyrosine kinase c-Met has been implicated in the progression of a variety of human cancers. In our previous study, we identified a hit compounds **1a** with the activity of 9279 nM. A fragment-based drug optimization was performed based on 1a (**Figure 5A**). According to the binding mode (**Figure 5B**), **1a** could form a hydrogen bond with the main chain of Met1160 at hinge residues. In the gate area, **1a** had hydrophobic interactions with, and formed a hydrogen bond with Asp1222 of DFG motif. For the back cleft, **1a** could form a hydrogen bond with Glu1127 of αC helix. However, the low potency of **1a** indicated that there was still room for the activity improvement.

**Figure 5.**
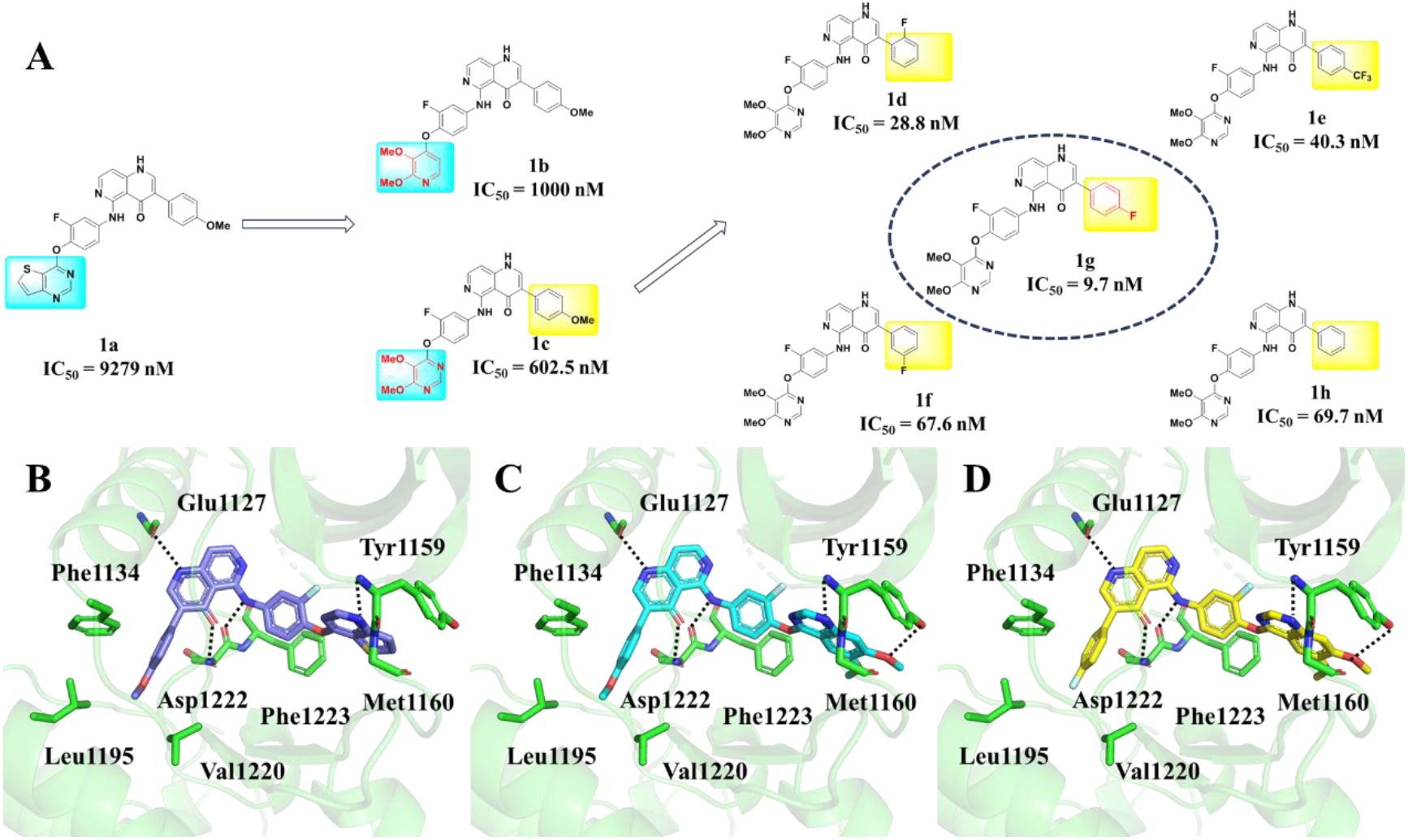
The discovery of potent c-Met inhibitor using KinaFrag. (A) The workflow of fragment-based optimization. **1a** (IC_50_ = 9279 nM) was used as the starting structure. The first step was to optimize the interaction with hinge residues. The most potent compound **1c** (IC_50_ = 602.5 nM) was selected for further evolution. The final result **1g** exhibited a IC_50_ value of 9.7 nM against c-Met. (B) The binding mode of **1a** with c-Met (PDB ID: 3F82). **1a** could form hydrogen bonds with Glu1127, Met1160, and Asp1222. (C) The binding mode of **1c** with c-Met (PDB ID: 3F82). **1c** could form hydrogen bonds with Glu1127, Tye1159, Met1160, and Asp1222. (B) The binding mode of **1g** with c-Met (PDB ID: 3F82). **1g** could form hydrogen bonds with Glu1127, Tye1159, Met1160, and Asp1222. The hydrogen bonds were shown in black dotted lines.

To optimize this compound, the modification was first aimed at the AP region. The interactions with hinge region play key roles for the activity of PKIs. From the docking result of **1a** with c-Met, we could find only one hydrogen bond was formed, which was less than most of PKIs and might affect the inhibitory activity. Therefore, the AP binding fragment dataset was used to conduct the fragment growing. According to the ΔG of new generated compounds (**Table S1**), compound **1b** and **1c** were synthesized, and exhibited a better bioactivity. As shown in the binding mode (**Figure 5C**), **1c** could form a hydrogen bond with the main chain of Met1160 at hinge region. Moreover, its methoxyl group could have another hydrogen bond with the side chain of Tyr1159. The increased hydrogen bond number was satisfied to the activity of **1b** (IC_50_ = 1000 nM) and **1c** (IC_50_ = 602.5 nM). The further optimization was performed to improve the interaction with the hydrophobic back cleft. The back cleft binding fragment library was employed to perform the fragment growing (**Table S2**). Five compounds were finally select to conduct the bioactivity evaluation. The most potent compound **1g** (IC_50_ = 9.7 nM), which had a ~1000-fold improvement in activity, could form the similar interactions with c-Met as **1c** (**Figure 5D**). In the back cleft, **1g** had hydrophobic interactions with the surrounding residues in the suitable distance. The in vitro experiment revealed that **1g** could inhibit the HGF-mediated autophosphoylation in MKN-45 cells with IC_50_ value of 47.3 nM. The successfully discovery of **1g** showed the reliability and efficiency of KinaFrag used for the fragment-based drug design.

## Conclusion

The understanding of kinase-fragment interaction and performing fragment-to-lead optimization are still significant challenges, which hindered the rapid discovery of selective PKIs. In this study, we construct a web source KinaFrag. KinaFrag contained 31464 disclosed kinase inhibitor fragments, which involved crystal structures and disclosed bioactive kinase inhibitors. The comprehensive fragment structure and kinase-fragment interaction information were significant to the design of PKIs. In addition, a fragment growing workflow was added in KinaFrag to explore the unknown kinase-fragment interaction space for novel inhibitor identification. We designed a more potent c-Met inhibitor with KinaFrag, and the *in vitro* activity was enhanced ~1000-fold compared with the hit compound. All users could view and download these resource freely. We hope KinaFrag could be helpful to the development of more promising PKIs for cancers and other diseases therapies.

## Key Points

Fragment-based drug discovery plays a vital role in the selective kinase inhibitor development. However, how to investigate kinase-fragment interaction and optimize fragment into hit compounds is still big challenges.

We have developed a new web resource for the fragment-based kinase inhibitor discovery, which represents the first web source for kinase-fragment interaction space exploration.

KinaFrag contained fragments as well as their binding information and multiple properties from protein kinase inhibitors with promising bioactivity, which could be used to investigate the known kinase-fragment interaction space for selective kinase inhibitor discovery.

Users could explore the unknown kinase-fragment interaction space and perform kinase inhibitor discovery via a fragment-growing strategy and fragments generated by KinaFrag.

KinaFrag can be used straightforward without need of registration.

## Acknowledgements

This work was supported by the National Key R&D Program (2018YFD0200100), the National Natural Science Foundation of China (21772059, 91853127, and 31960548). Program of Introducing Talents of Discipline to Universities of China (111 Program, D20023). Frontiers Science Center for Asymmetric Synthesis and Medicinal Molecules, Department of Education, Guizhou Province [Qianjiaohe KY number (2020)004].

## Author contributions

G.F.H. and G.F.Y designed the project. Z.Z.W. and F.W. collected the data. Z.Z.W. and F.W. developed the program of the database and fragment growing workflow. Z.Z.W., F.W. and X.X.S analyzed the results. G.F.Y., G.F.H., Z.Z.W. and F.W. wrote the manuscript.

## Conflicts of interest

There are no conflicts to declare.

## Notes

http://chemyang.ccnu.edu.cn/ccb/database/KinaFrag/index.php

### Competing Interest Statement

The authors have declared no competing interest.

